# The Chd1 chromatin remodeler shifts hexasomes unidirectionally

**DOI:** 10.1101/080010

**Authors:** Robert F. Levendosky, Anton Sabantsev, Sebastian Deindl, Gregory D. Bowman

## Abstract

Despite their canonical two-fold symmetry, nucleosomes in biological contexts are often asymmetric: functionalized with post-translational modifications (PTMs), substituted with histone variants, and even lacking H2A/H2B dimers. Here we show that the Widom 601 nucleosome positioning sequence can be used to produce hexasomes in a specific orientation on DNA, which provide a useful tool for interrogating chromatin enzymes and allow for the generation of precisely defined asymmetry in nucleosomes. Using this methodology, we demonstrate that the Chd1 chromatin remodeler requires H2A/H2B on the entry side for sliding, and thus, unlike the back-and-forth sliding observed for nucleosomes, Chd1 shifts hexasomes unidirectionally. Chd1 takes part in chromatin reorganization surrounding transcribing RNA polymerase II (Pol II), and using asymmetric nucleosomes we show that ubiquitin-conjugated H2B on the entry side stimulates nucleosome sliding by Chd1. We speculate that biased nucleosome and hexasome sliding due to asymmetry contributes to the packing of arrays observed in vivo.

## Introduction

As the repeating unit of chromatin, the nucleosome is the canvas upon which the epigenetic histone code is written. A fundamental characteristic of the histone code is the combinatorial diversity achieved from multiple marks, which may or may not reside on the same histone tail (Ruthenburg et al., 2007; Tee & Reinberg, 2014). Both through post-translational modifications (PTMs) and substitution of histone variants, additional chemical diversity arises from asymmetric modifications of nucleosomes. Since the nucleosome is pseudo-symmetric with two copies of each core histone (H2A, H2B, H3 and H4), asymmetry occurs when each copy possesses distinct epigenetic modifications. Recent advances have revealed asymmetry at the single nucleosome level (Rhee et al., 2014; Voigt et al., 2012), yet with challenges in synthesizing uniform populations of asymmetrically modified nucleosomes (Lechner et al., 2016; Liokatis et al., 2016), the biological significance of the vast majority of asymmetric marks remains unclear.

A dramatic example of asymmetry is the pairing of activating H3K4me3 and repressive H3K27me3 marks, known as bivalency (Voigt et al., 2013). Trimethylation of H3K27 is carried out by PRC2, and while H3K4me3 blocks modification of K27 on the same H3 tail, PRC2 can deposit H3K27me3 mark on the opposing H3 tail of the same nucleosome (Lechner et al., 2016; Voigt et al., 2012). In addition to generating nucleosomes with asymmetric H3K4me3/H3K27me3, PRC2 is also activated by the mark it deposits, with substrate preference for asymmetric nucleosomes containing one H3K27me3 (Lechner et al., 2016; Margueron et al., 2009). While recognition of asymmetric H3K27me3 is believed to be important for maintenance and spreading of heterochromatin, and the bivalent H3K4me3/H3K27me3 signature has been well established for stem cell identity, there is relatively little biological understanding for most other epigenetic marks that are prominently asymmetric. Genome-wide studies have revealed that the +1 nucleosome is strikingly asymmetric with regards to H3K9 acetylation, H2B ubiquitinylation, and residency of H2A.Z (Rhee et al., 2014). Asymmetric marks of the +1 nucleosome correlate with asymmetric localization of the RSC, INO80, and SWR1 chromatin remodelers (Ramachandran et al., 2015; Yen et al., 2012), and a major question is deciphering how these and other enzymes generate and read-out the asymmetric distribution of these marks.

Nucleosomes can also exhibit asymmetry with respect to histone content, with the lack of one H2A/H2B dimer defining the hexasome. The existence of hexasomes in vivo has been supported by ChIP-Exo and MNase-seq experiments (Rhee et al., 2014), and in vitro, hexasomes have been shown to be generated by the RSC remodeler with NAP1 histone chaperone (Kuryan et al., 2012) and also by RNA polymerase II (Pol II) transcribing through nucleosomes (Kireeva et al., 2002; Kireeva et al., 2005). Intriguingly, Pol II successfully transcribes through hexasomes oriented with the promoter-distal H2A/H2B dimer missing, but stalls in the absence of the promoter-proximal dimer (Kulaeva et al., 2009). Whether the orientation of hexasomes may affect other enzymes that act on chromatin has not previously been addressed.

Transcription requires local disruption and reassembly of nucleosomes, which is achieved by elongation factors, histone chaperones, and chromatin remodelers such as Chd1 and ISWI (Venkatesh & Workman, 2015). Chd1 and ISWI reposition nucleosomes into evenly spaced arrays, and are required for packing arrays of nucleosomes against the +1 nucleosome (Gkikopoulos et al., 2011; Lusser et al., 2005; Pointner et al., 2012; Tsukiyama et al., 1999). Although specific binding of H3K4me3 by the chromodomains of mouse Chd1 has been correlated with its localization to the promoter (Lin et al., 2011), Chd1 and ISWI remodelers have been shown to participate in resetting the chromatin barrier in coding regions after passage of Pol II, required for preventing cryptic transcription (Cheung et al., 2008; Pointner et al., 2012; Radman-Livaja et al., 2012; Smolle et al., 2012). Chd1 has been linked to elongating Pol II through interactions with the transcriptional elongation factors FACT and Spt4-Spt5, and with the Rtf1 subunit of the PAF complex (Kelley et al., 1999; Krogan et al., 2002; Simic et al., 2003). To aid passage of Pol II, the machinery that travels along with the transcription bubble alters local chromatin structure, yet it is not known how changes to nucleosomes might influence Chd1 or other chromatin remodelers. In addition to potentially generating hexasomes, passage of Pol II is also coupled to transient ubiquitinylation of H2B (Fleming et al., 2008; Xiao et al., 2005). Interestingly, the H2B-Ubiquitin (H2B-Ub) mark is required for FACT-assisted disruption of the chromatin barrier (Pavri et al., 2006). Chd1 has been shown to be required for high levels of transcription-coupled ubiquitinylation of H2B in vivo (Lee et al., 2012), yet a direct connection between Chd1 and transcriptionally altered nucleosomes has remained elusive.

In this work, we report the discovery that the Widom 601 nucleosome positioning sequence can generate oriented hexasomes, with the sole H2A/H2B positioned in a sequence-defined location. Using oriented hexasomes, we show that Chd1 requires H2A/H2B on the entry side for robust sliding and preferentially shifts hexasomes unidirectionally. Hexasomes can be transformed into nucleosomes upon addition of H2A/H2B dimers, and we demonstrate that oriented hexasomes are an ideal substrate for generating uniform populations of asymmetric nucleosomes with uniquely modified H2A/H2B dimers. We find that nucleosomes with an asymmetric H2B-Ub modification stimulate nucleosome sliding by Chd1, revealing an unexpected activating role for H2B-Ub in remodeling.

## Results

### The Widom 601 sequence allows for generation of oriented hexasomes

Since the nucleosome consists of two copies each of the four canonical histones – H2A, H2B, H3 and H4 – in vitro nucleosome reconstitutions that deviate from equi-molar histone stoichiometries can result in sub-nucleosomal products. Curiously, during the course of nucleosome reconstitutions by salt dialysis, we noticed that native PAGE migration of a smaller species changed depending on the location of flanking DNA. We use the strong Widom 601 positioning sequence (Lowary & Widom, 1998), with the X-601-Y naming convention, where X and Y refer to the number of base pairs flanking the core 145 bp 601 sequence. Consistently, we observed that the subspecies from 80-601-0 preps migrated faster than that of 0-601-80 preps (Figure 1A). The hexasome is a stable sub-nucleosomal particle lacking one of the two H2A/H2B dimers (Arimura et al., 2012; Kireeva et al., 2002; Mazurkiewicz et al., 2006), and we confirmed by SDS-PAGE analysis that the faster migrating species were in fact hexasomes (Figure 1B).

**Figure 1.**
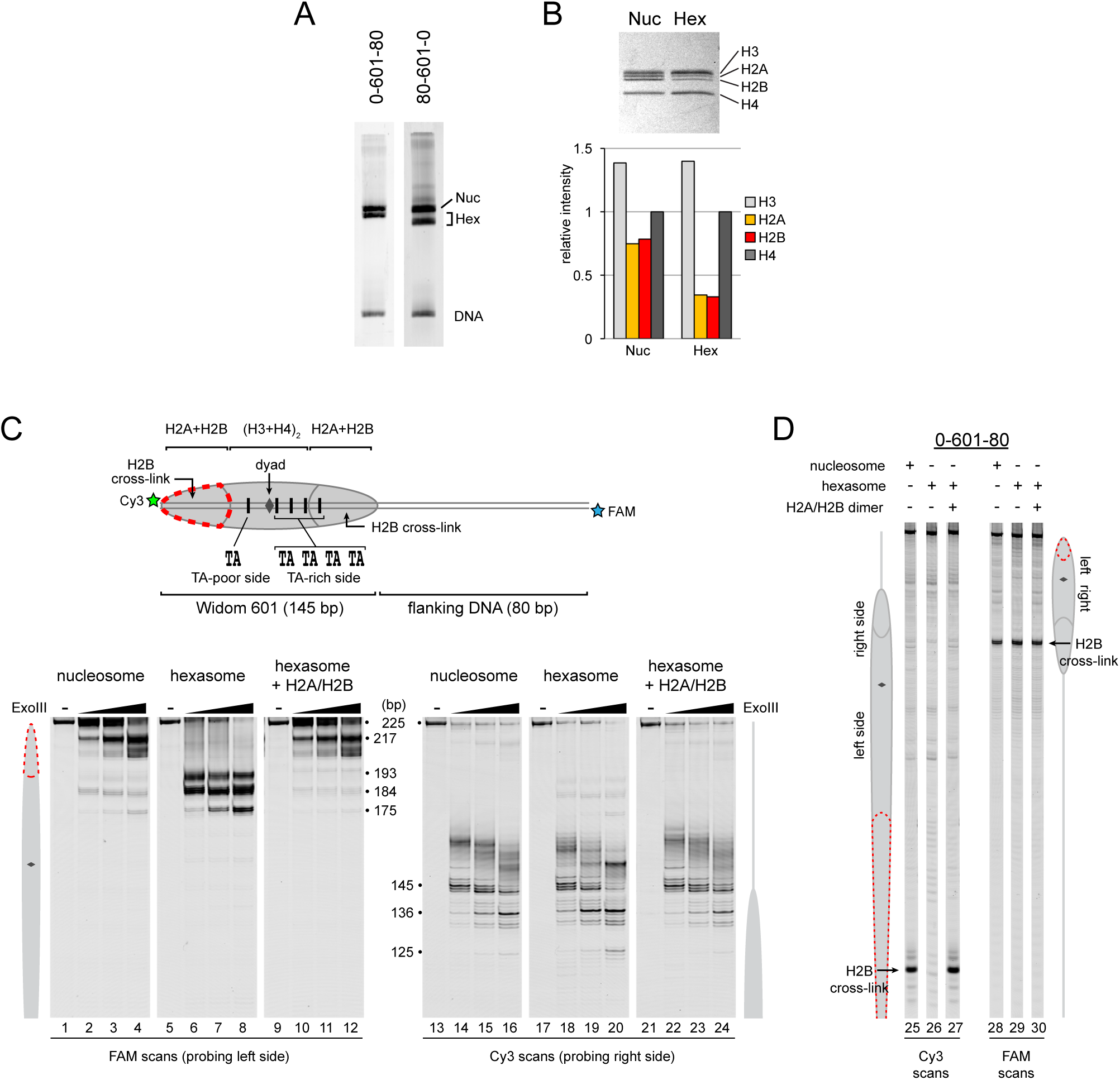
Oriented hexasomes can be generated using the Widom 601 sequence. **(A)** Hexasomes but not nucleosomes migrate differently by native PAGE when flanking DNA is on the left or right of the 601 sequence. These two gels, poured from the same solution, are representative of typical 0-N-80 and 80-N-0 reconstitutions. (**B**) Analysis of histone stoichiometry by SDS-PAGE confirms that hexasome species lacks one H2A/H2B dimer. The bar graph is a quantification of the gel shown, and band intensities were normalized to histone H4. The gel shown is representative of three similar experiments using different hexasome/nucleosome preparations. (**C**) Exonuclease III analysis demonstrates that hexasomes only retain one H2A/H2B dimer on the TA-rich side of the 601 sequence. Samples were incubated with 0, 10, 40, and 160 units of ExoIII and resolved on urea denaturing gels. Lanes 9-12 and 21-24 show addition of 200 nM H2A/H2B dimer to 100 nM hexasomes, which recovered nucleosome digestion patterns. The size (bp) of major products are indicated. These gels are representative of two independent experiments. (**D**) Histone mapping confirms unique the orientation of hexasomes on the Widom 601 and the generation of nucleosomes with H2A/H2B dimer addition. Related mapping experiments using wild type and modified hexasomes were performed four times with similar results. See also Figure 1 - figure supplement 1.

Nucleosomes migrate differently in native gels depending on whether flanking DNA is present only on one or both sides of the histone core (Eberharter et al., 2004; Pennings et al., 1991). With one H2A/H2B dimer missing, a hexasome would have an additional ~40 bp of DNA extending from the core at a more internal location. We reasoned that the differences in hexasome migration may therefore be due to a systematic loss of H2A/H2B from one side of the Widom 601 sequence. To test this idea, we probed the accessibility of DNA using Exonuclease III (Figure 1C). Nucleosomes digested with ExoIII showed the expected protection at the edge of the histone core, with preferential cleavage in ~10-11 nt increments (lanes 2-4 and 14-16). The 0-601-80 hexasome, in contrast, was digested more internally by ~30-40 nt on the 0 bp side, while showing full nucleosome protection on the 80 bp side (lanes 6-8 and 18-20). Relative to the orientation of the 601 sequence, the 80-601-0 hexasome showed an analogous pattern, with ~30-40 nt more extensive ExoIII digestion on the 80 bp side and similar protection on the 0 bp side compared to nucleosomes (Figure 1 - figure supplement 1A). Thus, the preferred location of the remaining H2A/H2B dimer in the hexasome was not influenced by flanking DNA, but instead was determined in a sequence-specific fashion based on the orientation of the Widom 601.

Previous work by several labs has revealed asymmetry in the Widom 601 sequence with respect to the strength of histone-DNA contacts. Single-molecule DNA unzipping experiments demonstrated that one side of the 601 forms more stable contacts with histones (Hall et al., 2009), and the asymmetry of the 601 was found to form a polar barrier to passage of RNA polymerase II (Bondarenko et al., 2006). One feature that has been pointed out as a key determinant of stable histone-DNA contacts are periodic TA dinucleotide steps (Lowary & Widom, 1998). The Widom 601 is notably asymmetric in TA steps on either side of the dyad where binding affinity is expected to be highest, with four TA steps on one side opposite a single TA step (Chua et al., 2012). Generation of symmetric derivatives of 601 has shown that the TA-rich side is much more salt stable than the TA-poor side (Chua et al., 2012), and single molecule experiments have found that the TA-poor side preferentially unwraps under force (Ngo et al., 2015). We orient the Widom 601 with the TA-rich side on the right, which means that the side lacking the H2A/H2B dimer in hexasomes corresponds with the TA-poor side of the 601 sequence.

Others have shown that hexasomes can generate nucleosome-like products upon addition of H2A/H2B dimer (Kireeva et al., 2002). To investigate this step-wise method of generating nucleosomes, we incubated hexasomes with a 2-fold molar excess of H2A/H2B dimer and monitored ExoIII digestion. As shown in Figure 1C, addition of H2A/H2B dimer yielded a protection pattern indistinguishable from nucleosomes (lanes 10-12). Therefore, even in the absence of histone chaperones or elevated salt, addition of H2A/H2B dimer to hexasomes was sufficient for recovering nucleosome-like protection patterns.

As an alternative method for characterizing hexasomes and nucleosomes generated from H2A/H2B dimer addition, we used histone mapping. By labeling a single cysteine variant of H2B (S53C) with photo-reactive 4-azidophenacyl bromide (APB), UV-induced cross-linking can reveal the precise position of the H2A/H2B dimer on nucleosomal DNA (Abbott et al., 2005; Kassabov et al., 2002). In nucleosomes, each H2B cross-links to only one DNA strand, and therefore doubly-labeled fluorescent DNA is needed to report on both sides of the 601 sequence. In agreement with ExoIII experiments, H2B cross-linking for hexasomes was virtually absent on the TA-poor side of the 601, whereas cross-linking on the TA-rich side was equivalent for hexasomes and nucleosomes (Figure 1D; compare lane 25 with 26 and 28 with 29). Strikingly, addition of H2A/H2B fully recovered the H2B cross-link on the TA-poor side (compare lanes 26 and 27), demonstrating that dimer addition generates correctly organized nucleosomes that are indistinguishable from those obtained by salt dialysis reconstitution. Similar results were obtained with 80-601-0 hexasomes (Figure 1 - figure supplement 1B), reinforcing the conclusion that salt dialysis deposits limiting H2A/H2B on the TA-rich side of the Widom 601.

### Chd1 requires the entry-side H2A/H2B dimer for robust sliding

Given the strong sequence-defined placement of limiting H2A/H2B dimer, we refer to hexasomes produced by the Widom 601 as “oriented hexasomes.” By having a defined orientation on DNA, these hexasomes offer a unique tool for probing requirements of nucleosome-interacting enzymes. Despite the two-fold pseudo-symmetry of the nucleosome, factors binding off the central dyad axis encounter the two halves of the nucleosome at distinct distances and orientations. The two H2A/H2B dimers and the DNA they coordinate are therefore likely to play unequal roles in nucleosome recognition and enzyme regulation. Chromatin remodelers such as Chd1 shift DNA past the histone core by acting at an internal DNA site called superhelical location 2 (SHL2), located ~20 bp from the dyad (McKnight et al., 2011; Saha et al., 2005; Schwanbeck et al., 2004; Zofall et al., 2006). Relative to the SHL2 site of DNA translocation, one H2A/H2B dimer binds to DNA being shifted onto the nucleosome, and is therefore considered to be on the entry side, whereas the other H2A/H2B binds DNA about to be shifted off the histone core, and thus is on the exit side. In vitro, Chd1 slides mononucleosomes away from DNA ends (McKnight et al., 2011; Stockdale et al., 2006). By using end-positioned nucleosomes, we can restrict the direction of sliding, thereby defining H2A/H2B adjacent to the long flanking DNA as the entry-side dimer. Since the placement of the single H2A/H2B dimer relative to 601 is maintained regardless of flanking DNA (Figure 1; Figure 1 - figure supplement 1A), we can generate hexasomes with the H2A/H2B dimer on either the entry or exit side. This unique tool therefore allows us to determine the extent that Chd1 relies on either the entry- or exit-side H2A/H2B dimer.

As a standard technique for visualizing repositioning of mononucleosomes along DNA, we first investigated movement of oriented hexasomes using native PAGE (Figure 2A). We used 80-601-0 and 0-601-80 constructs described above, which lacked one of the H2A/H2B dimers on either the entry side (80-601-0) or exit side (0-601-80). Upon addition of Chd1 and ATP, nucleosomes mobility decreased, signifying movement away from DNA ends (lanes 1-8). In contrast, under the same conditions the hexasomes failed to show any clear changes in migration patterns (lanes 9-16).

**Figure 2.**
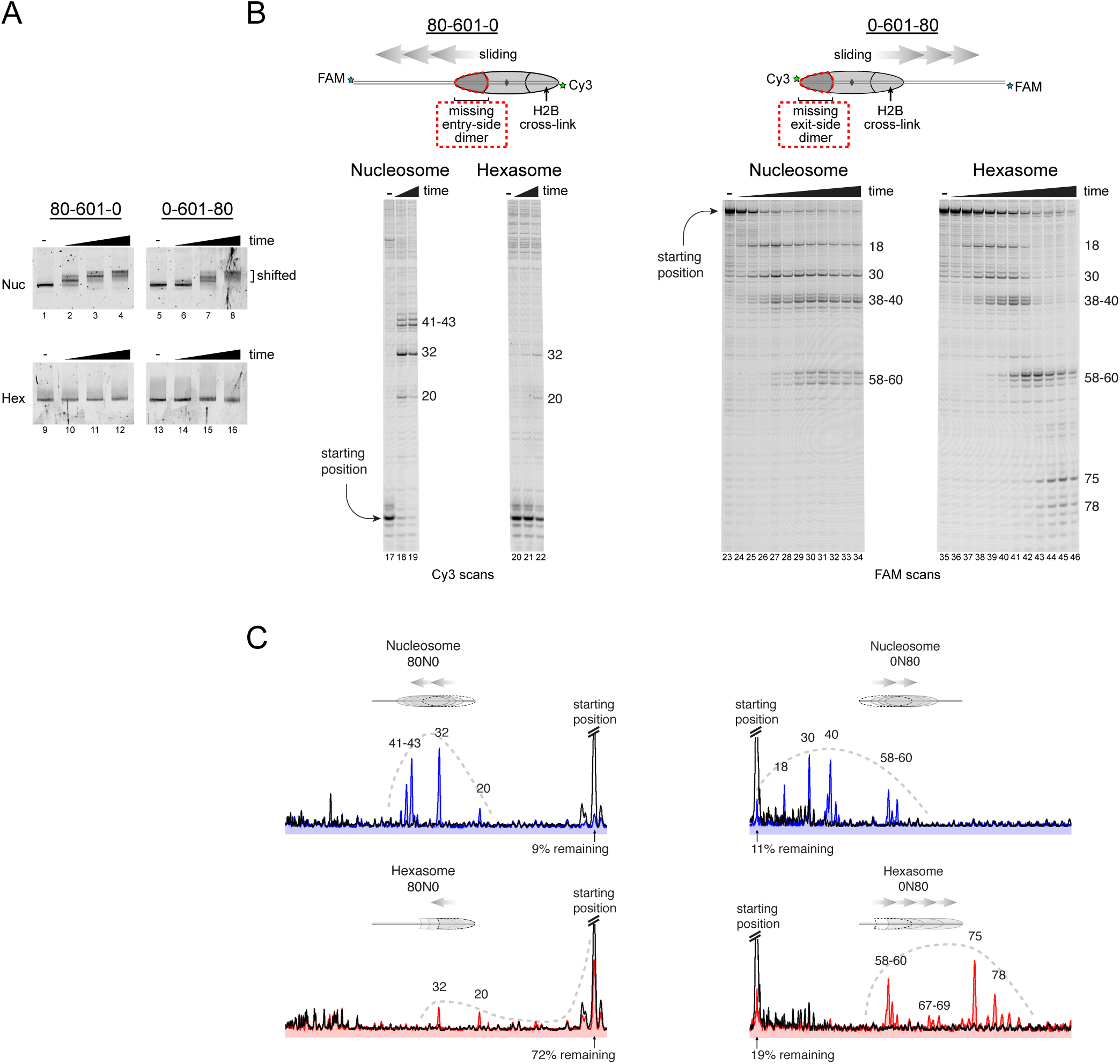
Chd1 requires the H2A/H2B dimer on the entry side for robust remodeling. (A) Nucleosome and hexasome sliding reactions analyzed by native PAGE. Nucleosomes or hexasomes (150 nM) were incubated with yeast Chd1 (50 nM) and 2.5 mM ATP and reactions resolved for 0, 0.5, 5, and 50 min time-points. Shown is a representative of four independent experiments. (**B**) Nucleosome and hexasome sliding reactions, visualized through histone mapping. For hexasome and nucleosome 80-601-0 constructs (left), sliding reactions were monitored after incubation with 50 nM Chd1 for 0, 1, and 64 min. For 0-601-80 constructs (right), time points were 0, 0.25, 0.5, 1, 2, 4, 8, 16, 32, and 64 min. Sliding experiments for 0-601-80 and 80-601-0 were each examined six or more times with similar results. (**C**) Comparison of intensity profiles for histone mapping reactions shown in (B). Samples before ATP addition (0 min) are black, nucleosome sliding reactions after 64 min are blue, and hexasome sliding reactions after 64 min are red.

Since the loss of one H2A/H2B dimer in the hexasome dramatically alters the location and thus the angle at which DNA extends away from the histone core, we considered the possibility that native PAGE may not accurately report on changes in hexasome positioning. We therefore investigated the ability of Chd1 to remodel hexasomes using histone mapping (Figure 2B). In agreement with previous work (Patel et al., 2013; Stockdale et al., 2006), Chd1 shifted end-positioned mononucleosomes to more central locations on DNA, with the majority of nucleosomes repositioned ~20 to ~60 bp from their starting locations (lanes 17-19, 23-34). For hexasomes, however, the ability of Chd1 to reposition was strongly dependent on the location of the single H2A/H2B dimer. The 80-601-0 hexasomes, which lacked the entry-side H2A/H2B dimer, failed to show robust repositioning, with the majority of products remaining at the starting position (lanes 20-22). In marked contrast, the 0-601-80 hexasomes shifted robustly onto flanking DNA, demonstrating that Chd1 activity can be supported by an H2A/H2B dimer on only the entry side (lanes 35-46). Interestingly, instead of generating more centrally positioned products, Chd1 shifted 0-601-80 hexasomes to the opposite end of DNA, farther than observed for nucleosomes (Figure 2B,C). This biased movement of 0-601-80 is consistent with the poor sliding of 80-601-0 hexasomes, which lack entry-side H2A/H2B, and indicate that Chd1 preferentially slides hexasomes toward the side containing the H2A/H2B dimer. Thus, Chd1 can reposition hexasomes, but the requirement for entry-side H2A/H2B yields a strong directional bias for hexasomes that contrasts with the back-and-forth sliding observed for nucleosomes.

### Oriented hexasomes allow for precisely designed asymmetric nucleosomes

The discovery of oriented hexasomes opens up a simple means for producing asymmetric nucleosomes, where unique modifications in the two H2A/H2B dimers can be directed to specific sides of the nucleosome. One powerful technique that can benefit from generating asymmetric nucleosomes is single molecule FRET (smFRET). Though many variations are possible, fluorescent dye labeling of nucleosomes commonly involves both histones and DNA, which allows for detection of DNA unwrapping and DNA translocation relative to the histone core (Li & Widom, 2004; Yang et al., 2006; Blosser et al., 2009). The FRET signal, however, can be complicated by the two-fold symmetry of the nucleosome, since dyes at the two related histone positions are typically not equidistant from the DNA-tethered dye, and therefore lead to a mixture of FRET levels (Deindl et al., 2013). A standard solution to this issue has been to dilute the labeled histone with an excess of unlabeled histone during nucleosome reconstitution, and select out the desired FRET signal from a single donor/acceptor pair. We expected that the unique placement of a single H2A/H2B dimer relative to the DNA sequence should allow us to generate nucleosomes with a single, uniform FRET pair.

To examine this idea, we labeled H2A-T120C with Cy3-maleimide and generated 80-601-3 hexasomes and nucleosomes containing a DNA-tethered Cy5 dye on the 3 bp side. As previously described (Deindl et al., 2013), the nucleosomes gave rise to two major FRET populations corresponding to single Cy3 dyes on the distal or proximal H2A (Figure 3A). A mid-FRET population is expected between these two major species, where nucleosomes contain Cy3 on both copies of H2A copies. Here, due to extensive dilution of labeled H2A-Cy3, that population was minimal. In contrast, the oriented hexasomes yielded a single, high-FRET population as expected for the H2A/H2B dimer located on the exit side, proximal to the DNA label (Figure 3B). To see whether these hexasomes would behave as nucleosomes upon H2A/H2B dimer addition, we incubated these samples with Chd1 and ATP to stimulate nucleosome sliding. After a 10 min incubation, all nucleosomes had shifted to a low FRET state, as expected for a ≥ 20 bp shift of the histone core away from the labeled DNA end. Hexasomes, in contrast, maintained a significant population of high-FRET species after incubation, consistent with the poor movement observed by histone mapping in the absence of entry-side H2A/H2B dimer. Addition of H2A/H2B dimer to hexasomes did not significantly alter the starting high-FRET population, yet incubation with Chd1 and ATP yielded a low-FRET profile similar to that observed for nucleosomes (Figure 3C). These results show that oriented hexasomes offer a defined methodology for producing uniformly labeled nucleosomes that should benefit smFRET experiments.

**Figure 3.**
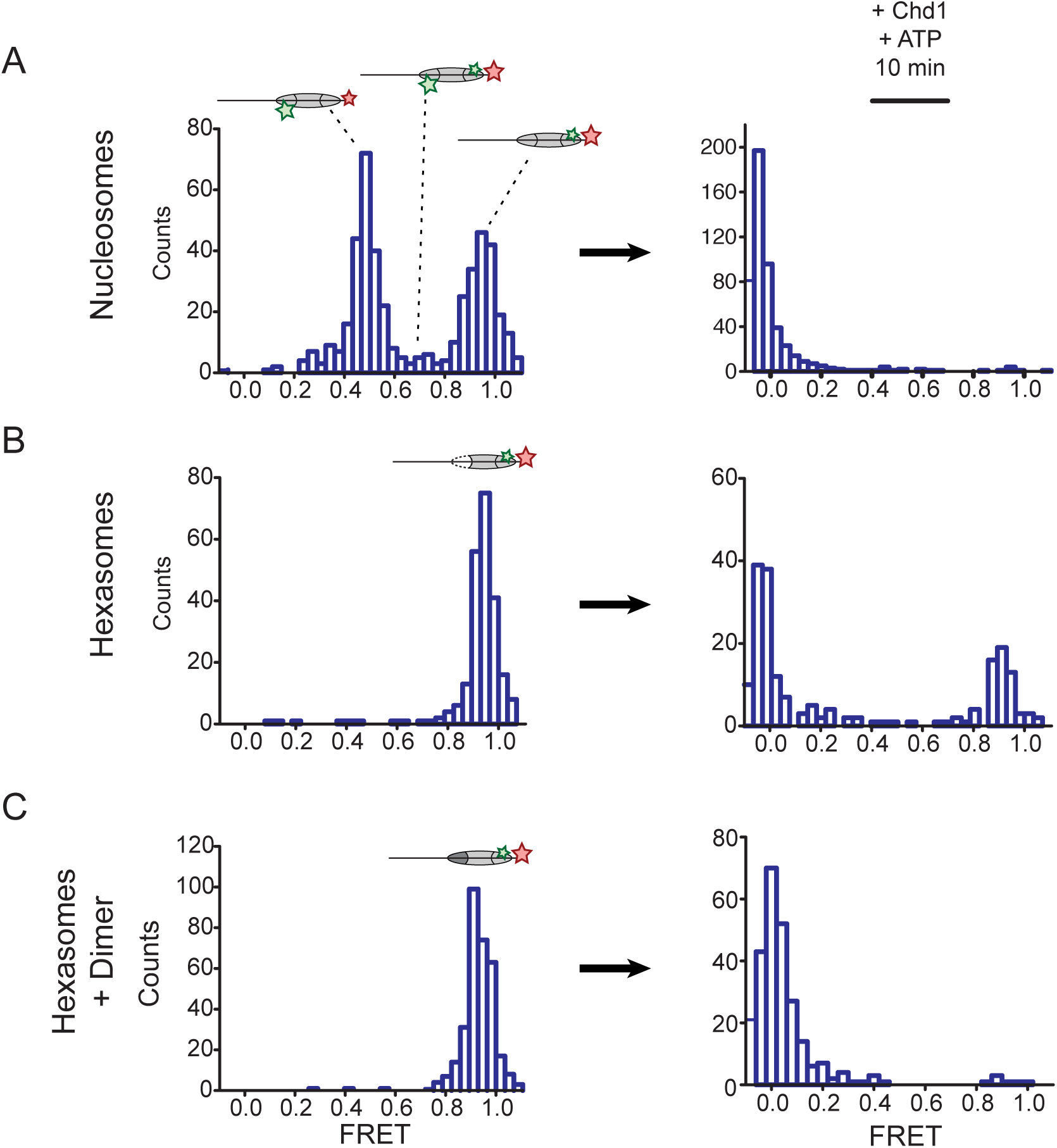
Oriented hexasomes allow targeted placement of modified H2A/H2B dimers on the nucleosome. **(A)** Analysis of dual labeled nucleosomes (H2A-Cy3 and DNA-Cy5) by single-molecule FRET (smFRET) reveals multiple species prior to nucleosome sliding by Chd1. **(B)** Oriented hexasomes (H2A-Cy3 and DNA-Cy5) made with an 80-601-3 uniformly show one dye pair that yields high FRET. Right panel shows relatively poor mobilization of hexasomes by Chd1. (**C**) Addition of unlabeled H2A/H2B dimer to the labeled 80-601-0 hexasomes yields asymmetric nucleosomes, only possessing the high FRET dye pair.

### Quantification of remodeling activity using nucleosomes produced from hexasomes plus H2A/H2B dimer

The experiments presented above suggested that the addition of H2A/H2B dimer to hexasomes was sufficient for producing properly organized nucleosomes. For measuring kinetics of chromatin remodeling, however, we were concerned about two possible complications associated with H2A/H2B dimer addition: too little dimer would allow a significant fraction of hexasome to remain in the reaction, which might compete with the nucleosome, whereas too much dimer could create off-products that might interfere with nucleosome sliding. To evaluate these potential effects of H2A/H2B dimer addition, we monitored nucleosome sliding reactions using an assay based on static quenching of fluorescence (SQOF). We found that fluorescence quenching of two Cy3 dyes, placed on histone H2A(T120C) and the “zero” DNA end, is reduced when the zero-end (exit side) is shifted farther away from the histone core. Using this labeling scheme, we monitored changes in fluorescence intensity of 80-601-0 hexasomes, possessing Cy3-H2A on the exit dimer, upon the addition of unlabeled H2A/H2B dimer, Chd1, and ATP (Figure 4A). Relative to the constant hexasome concentration used (10 nM), addition of unlabeled H2A/H2B dimer stimulated sliding at all concentrations, from undersaturating (4 nM) to saturating (24 nM). The total change in fluorescence intensity increased with dimer concentration, consistent with only the fraction of hexasomes converted to nucleosomes being readily shifted by Chd1. A maximum change in fluorescence was observed with a 1.6-fold molar ratio of dimer to hexasome (Figure 4B), which is in agreement with others who have reported requiring ~2-fold molar equivalent of dimer to convert all hexasome to nucleosome (Kireeva et al., 2002).

**Figure 4.**
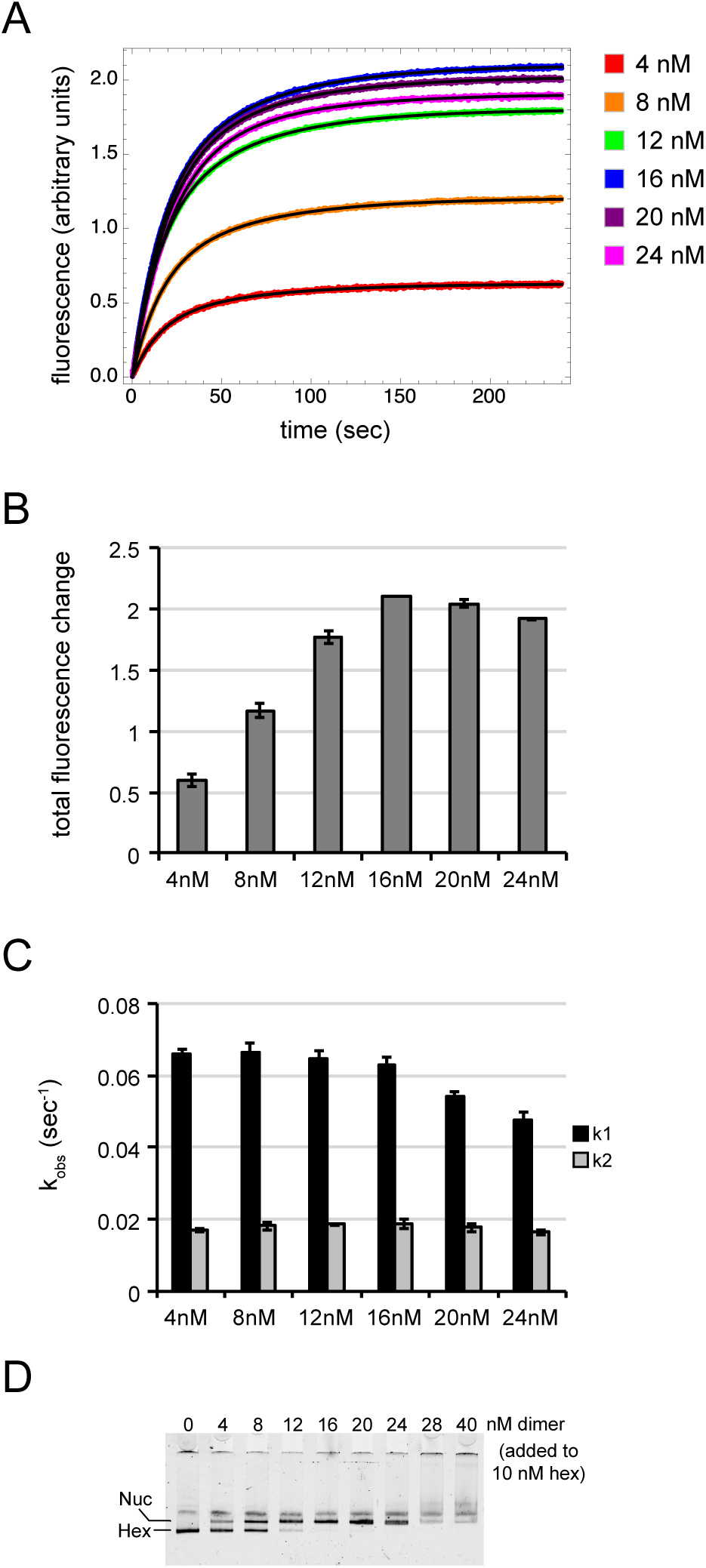
With subsaturating H2A/H2B dimer addition, rates of nucleosome sliding by Chd1 are not sensitive to nucleosome:hexasome ratios. (**A**) Nucleosome sliding monitored by Cy3-Cy3 fluorescence quenching. The legend shows the amount of H2A/H2B dimer for each reaction, which contained 10 nM 80-601-0 hexasome. Each trace is an average of 4 or more injections from the same stopped flow experiment. The black curves show double exponential fits to the data. (**B**) Graph of overall intensity changes at each H2A/H2B dimer concentration added to hexasomes, with higher intensity reflecting a greater proportion of nucleosomes that can be shifted. Error bars indicate the range from two independent sets of experiments. (**C**) Graph of sliding rates for stopped flow H2A/H2B dimer addition. Rates (k1 and k2) were determined from a double exponential fit to the data. Error bars indicate the range from two independent sets of experiments. (**D**) Native PAGE visualization of nucleosomes generated by addition of H2A/H2B dimer to hexasomes. Shown is a representative of ten similar titrations performed using wild-type or modified H2A/H2B dimers.

Interestingly, the reaction rates were maximal and remained constant over a wide range of H2A/H2B concentrations, from subsaturating up to the 1.6-fold molar equivalent that yielded the maximum change in fluorescence intensity (Figure 4C). Thus, under the conditions used here, the presence of hexasome due to limited H2A/H2B dimer addition did not influence rates for Chd1 remodeling. Beyond this saturating amount, both the rates and range of fluorescence intensity decreased. These reductions likely resulted from improper H2A/H2B deposition on flanking DNA that interferes with Chd1 action. We also monitored nucleosome formation by native PAGE, which showed a dimer-mediated shift of the hexasome species to nucleosomes and aggregation with excessive H2A/H2B (Figure 4D). As the most consistent rates were observed below the two-fold molar equivalent, we used only a slight molar excess of dimer for the remainder of our dimer addition experiments (10 nM hexasome plus 12 nM H2A/H2B).

### Chd1 is specifically stimulated by ubiquitinylated H2B on the entry-side dimer

The ability to separately add each H2A/H2B via oriented hexasomes offers a straightforward methodology to test whether particular enzymes are influenced by histone modifications on one or both sides of the nucleosome. Based on the close ties of both Chd1 and H2B ubiquitinylation with transcribing Pol II (Kelley et al., 1999; Krogan et al., 2002; Simic et al., 2003; Xiao et al., 2005), we were interested to determine whether Chd1 might be differentially influenced by a ubiquitin modification on one or both sides of the nucleosome.

We used oriented hexasomes to produce nucleosomes with the four combinations of modified and unmodified H2A/H2B dimers. To assemble nucleosomes with different placements of ubiquitin, we first generated 80-601-0 hexasomes with either unlabeled (Wt) or ubiquitinylated (Ub) H2B on the exit dimer. To these hexasomes, H2A/H2B dimer (Wt or Ub) was then added to each to produce the four combinations of Wt/Ub nucleosomes. Since ubiquitin significantly alters the sizes and shapes of hexasomes and nucleosomes, analysis by native gel clearly demonstrates that each reaction possesses a unique nucleosome with the desired placement of modified and unmodified H2A/H2B dimers (Figure 5A).

**Figure5.**
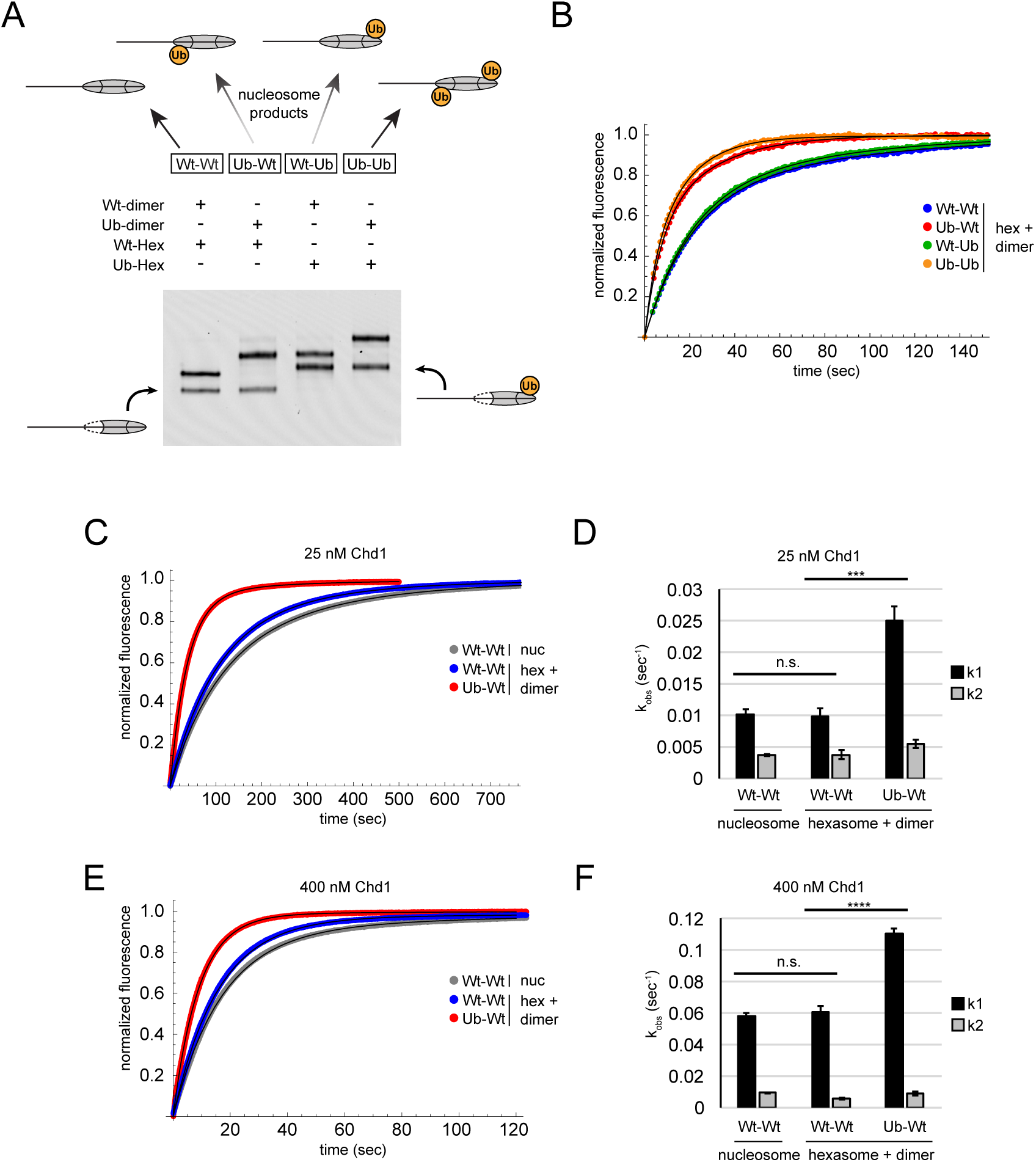
Entry-side H2B-Ubiquitin stimulates nucleosome sliding by Chd1. (**A**) Generation of symmetric and asymmetric nucleosomes with site-specific placement of H2B-Ubiquitin. Nucleosomes were formed from H2A/H2B dimer (12 nM) addition to 80-601-0 hexasomes (10 nM). Hexasomes and H2A/H2B dimer contained either unmodified (Wt) or ubiquitinylated (Ub) H2B as indicated, and resulting nucleosome and hexasome species were visualized by native PAGE. Shown is a representative from six independent dimer addition experiments. (**B**) Comparison of remodeling reactions with subsaturating (25 nM) Chd1, using hexasomes (10 nM) and H2A/H2B dimers (12 nM) containing unmodified or Ub-conjugated H2B. Shown are progress curves for remodeling reactions monitored on a fluorimeter using a Cy3-Cy3 pair. Black traces represent fits to the data. Progress curves are representative of two independent experiments. (**C**) Representative progress curves from three independent experiments of nucleosome sliding reactions monitored by stopped flow using Cy3B-Dabcyl at subsaturating (25 nM) Chd1. Black traces represent fits to the data. **(D)** Comparison of sliding rates monitored with Cy3B-Dabcyl at subsaturating Chd1 (25 nM). **(E)** Representative progress curves from three independent experiments of nucleosome sliding reactions monitored by stopped flow using Cy3B-Dabcyl at saturating (400 nM) Chd1. Black traces represent fits to the data. (**F**) Comparison of sliding rates monitored with Cy3B-Dabcyl at saturating Chd1 (400 nM). Error bars in x(D) and (F) show standard deviations from three independent experiments. *** P-value <0.0006; **** P-value <0.0001; n.s., not significant. See also Figure 5-figure supplement 1.

With these four nucleosome species, sliding reactions monitored by Cy3-Cy3 SQOF were carried out to determine whether H2A-Ub affected Chd1 activities. Despite the significant size of ubiquitin and potential to block access to H2A/H2B, the presence of this modification did not impede nucleosome sliding by Chd1. In fact, the two nucleosomes containing entry-side H2B-Ub (Ub-Wt and Ub-Ub) yielded faster rates than nucleosomes with unmodified entry-side H2B (Figure 5B). These results reinforce the finding that Chd1 activity is sensitive to the entry-side dimer, and reveal that Chd1 is stimulated by H2B-Ub.

During the course of these experiments, we found that binding of Chd1 to nucleosomes contributed to changes in Cy3 fluorescence due to protein induced fluorescence enhancement (PIFE). To determine the extent that the observed stimulation with H2B-Ub resulted from nucleosome sliding, we switched to Cy3B, a dye that does not allow for PIFE (Hwang et al., 2011), in combination with the quencher Dabcyl, which should exclusively report on nucleosome movement. Using the Cy3B-Dabcyl pair, we measured the rates of Chd1 sliding for nucleosomes with and without H2B-Ub on the entry side, generated from oriented hexasomes, as well as for unmodified nucleosomes purified from salt dialysis reconstitution. In agreement with previous experiments showing a preferential stimulation of Chd1, nucleosomes containing entry-side H2B-Ub consistently showed faster rates of nucleosome sliding (Figure 5C,D). For nucleosomes containing only unmodified H2B on entry and exit sides, no significant differences were observed in Chd1 sliding rates whether nucleosomes were produced from H2A/H2B addition to hexasomes or traditional generation via salt dialysis.

Since sliding experiments were performed using subsaturating (25 nM) concentrations of Chd1, it was unclear whether stimulation was due to a K_M_ or k_cat_ effect. To address this question, sliding experiments were repeated using a saturating (400 nM) concentration of Chd1 (Figure 5 - figure supplement 1). Interestingly, nucleosomes containing entry-side H2B-Ub were still shifted significantly faster with saturating Chd1, indicating that H2B-Ub did not merely improve Chd1 binding but increased catalytic efficiency (Figure 5E,F). The presence of the ubiquitin moiety could accomplish this either by helping to retain Chd1 on the nucleosome thereby increasing processivity/productivity from one enzyme-binding event, or by predisposing Chd1 to adopt an active conformation on the nucleosome.

## Discussion

Here we show how the Widom 601 can be used to generate oriented hexasomes, a unique and powerful tool for studying chromatin-interacting factors. As the highest affinity and most widely used nucleosome positioning sequence, the Widom 601 is well known for its marked asymmetry in strength of histone-DNA contacts. This asymmetry correlates with the distribution of TA dinucleotide steps where the minor groove contacts the histone core (Chua et al., 2012). The TA-rich side of the 601 has been shown to be more flexible and wrap more tightly (Chua et al., 2012; Hall et al., 2009; Ngo et al., 2015), and our results indicate that the TA-rich side of the 601 also has a higher affinity for the H2A/H2B dimer. We show that nucleosome reconstitution with limiting H2A/H2B dimer yields oriented hexasomes, with the single H2A/H2B dimer positioned in a sequence-specific fashion (Figure 1).

For chromatin remodelers, each H2A/H2B dimer is in a unique position, engaging with DNA either entering or exiting the nucleosome. Using oriented hexasomes, we show that Chd1 sliding activity strongly depends on the entry-side H2A/H2B dimer (Figure 2).

Nucleosome sliding requires engagement of the remodeler ATPase motor at one of the two internal SHL2 sites on the nucleosome (McKnight et al., 2011; Saha et al., 2005; Schwanbeck et al., 2004; Zofall et al., 2006). Like other remodelers, the ATPase motor of Chd1 shifts DNA toward the dyad, which means that the SHL2 site where the motor acts is adjacent to the entry-side H2A/H2B dimer. On hexasomes, Chd1 sliding appears restricted to the SHL2 adjacent to the sole H2A/H2B, resulting in virtually unidirectional movement (Figure 2B). These results reveal the importance of the entry-side H2A/H2B dimer for Chd1 sliding, which may arise from a critical epitope within H2A/H2B or a requirement for nucleosomal wrapping of entry-side DNA. Although experiments performed here were limited to the Chd1 remodeler, we expect that the biochemically similar ISWI remodelers, which slide nucleosomes directionally away from bound transcription factors and generate evenly spaced nucleosome arrays (Kang et al., 2002; Li et al., 2015; Lusser et al., 2005), should also exhibit a strong directional bias in sliding hexasomes.

The directional sliding of hexasomes by Chd1, which contrasts with the back-and-forth movement typical for nucleosomes, likely influences chromatin organization in vivo. In vitro, both transcription through nucleosomes by Pol II and remodeling by RSC along with the NAP1 chaperone have been shown to generate hexasomes (Kireeva et al., 2002). Although histone chaperones such as FACT would be expected to replace the missing H2A/H2B dimer during transcription, the passage of Pol II has been shown to specifically displace the H2A/H2B dimer distal to the promoter (Hsieh et al., 2013; Kulaeva et al., 2009), which would orient hexasomes for biased sliding toward the 5’ end. One speculative idea is that nucleosome array packing against the +1 nucleosome, which is dependent on Chd1 and ISWI chromatin remodelers (Gkikopoulos et al., 2011; Zhang et al., 2011), would be favored by directional hexasome sliding (Figure 6). Even with relatively few or transient hexasomes, directional sliding toward the transcriptional start site would be expected to corral upstream nucleosomes, similar to the phasing of nucleosome arrays against transcription factor-targeted nucleosomes (McKnight et al., 2016; Wiechens et al., 2016). Consistent with this idea, it has been shown that inactivation of Pol II relaxes nucleosome packing in coding regions, resulting in a nucleosome drift of ~10 bp toward the 3’ end of yeast genes (Weiner et al., 2010).

**Figure 6.**
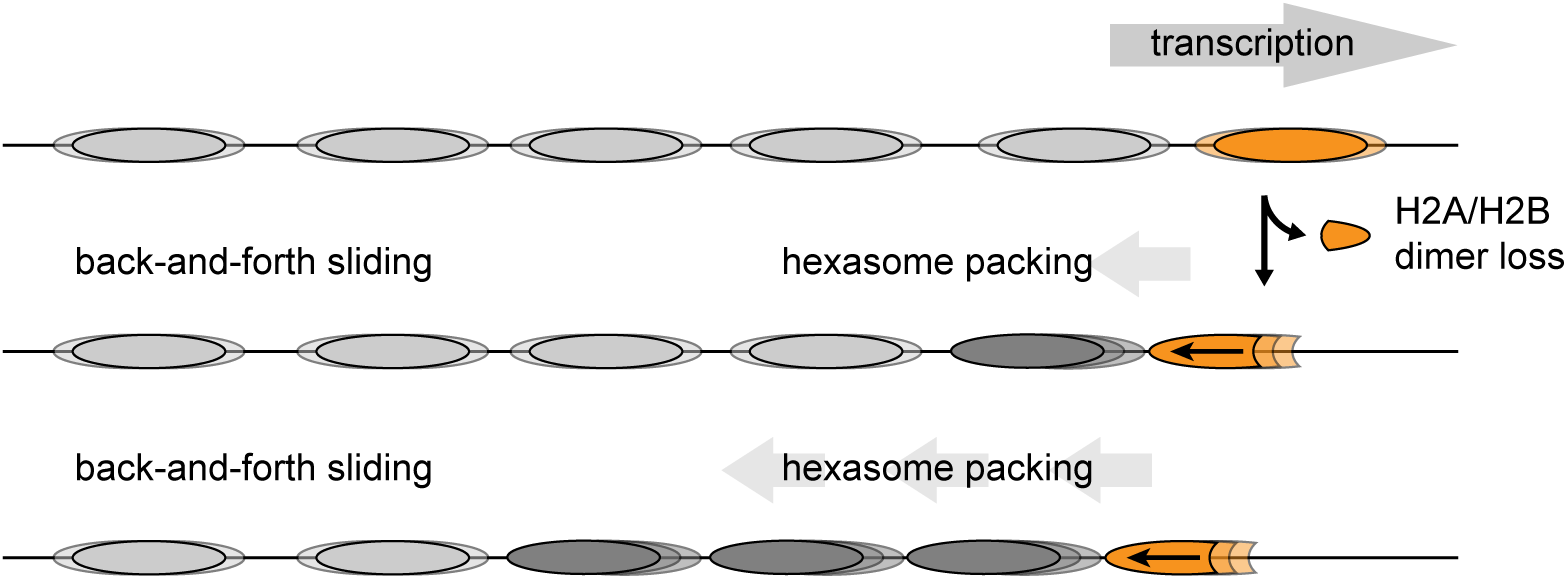
Model for nucleosome packing by oriented hexasomes. As others have shown, transcription by Pol II through nucleosomes is facilitated by removal of the promoter-distal H2A/H2B dimer (Kulaeva et al., 2009). Our results indicate that Chd1 would slide a hexasome of this orientation upstream. We propose that one or more hexasomes would corral intervening nucleosomes toward the promoter. Alternately, if every transcribed nucleosome were briefly converted to a hexasome, unidirectionally sliding of each hexasome would maintain tight nucleosome packing.

Our work also demonstrates how hexasomes made using the Widom 601 are a useful tool for generating specifically oriented asymmetric nucleosomes. To extract meaningful information on nucleosome dynamics, single molecule FRET experiments require a single, specifically positioned donor-acceptor pair (Blosser et al., 2009; Deindl et al., 2013; Ngo et al., 2015).

However, due to the pseudo two-fold symmetry of the nucleosome, labeling any of the histones with a FRET fluorophore (donor or acceptor) typically yields three different labeling configurations (Deindl et al., 2013). The presence of multiple FRET pairs, each with distinct and potentially closely spaced FRET values substantially complicates such analyses and at the same time limits throughput by decreasing the population of nucleosomes with the desired labeling configuration. Here, we show how oriented hexasomes can yield a homogeneously labeled population (Figure 3) that greatly facilitates smFRET experiments.

We furthermore demonstrate the utility of oriented hexasomes for generating asymmetric nucleosomes containing a uniquely positioned H2A/H2B PTM (>Figure 5A).

This method complements a recent procedure described for generating nucleosomes with distinct H3 tails (Lechner et al., 2016), and has the added advantage that the standard nucleosome reconstitution is sufficient for making oriented hexasomes, which can then be readily transformed into asymmetric nucleosomes with H2A/H2B dimer addition. Given the prevalence of asymmetric histone modifications and variants, this methodology should aid further investigation into how spatial cues contribute to the histone code, especially when used in conjunction with recent advances in detecting modifications on asymmetric nucleosome substrates (Liokatis et al., 2016).

Using this methodology, we demonstrate that ubiquitin-modified H2B stimulates Chd1, but only when present on the entry-side dimer (Figure 5). Unexpectedly, the stimulatory effect of ubiquitin was observed even with saturating remodeler concentrations, indicating that it is not merely the result of improved Chd1 binding. While we favor the idea of a direct interaction, where ubiquitin stabilizes an active conformation of Chd1 on the nucleosome, it is also possible that H2B-Ub alters the structure or dynamics of the nucleosome itself. Interestingly, H2B-Ub is required for maximum stimulation of Pol II transcription by FACT, potentially through aiding displacement of H2A/H2B (Pavri et al., 2006).

The stimulatory effect of ubiquitin raises new questions regarding how histone modifications bias chromatin remodelers, and more broadly, the discovery of oriented hexasomes should be a valuable tool for deepening our understanding of chromatin biology.

## Methods

### Protein production and modifications

Expression and purification of proteins used in this study were carried out as previously described for a truncated form of *S. cerevisiae* Chd1 (residues 118-1274) (Hauk et al., 2010; Patel et al., 2011), *Xenopus laevis* histones (Luger et al., 1999), and the ubiquitin variant G120C (Long et al., 2014). The ubiquitin sequence used in this work was identical to human and *Xenopus*, which is 96% identical to *S. cerevisiae* ubiquitin.

Conjugation of H2B and ubiquitin (H2B-Ub) was carried out essentially as described (Long et al., 2014) to produce a nonhydrolyzable H2B-Ub mimic. Briefly, based on the concentration of reduced cysteines using Ellman’s reagent, H2B(K120C) and His-tagged Ubiquitin (G76C) were combined at a 2:1 ratio at a protein concentration of ~ 10 mg・ml^−1^ in denaturing conditions. TCEP (5 mM C_f_) was added and incubated for 30 minutes. Crosslinking of the two proteins was carried out by adding 0.1 M 1,3 dichloroacetone to a final amount equal to half the total moles of reduced cysteines, and then quenching with 2-mercaptoethanol after 45 minutes. Un-crosslinked histones were removed using nickel affinity purification under denaturing conditions. The amount of crosslinked H2B-Ub was estimated from SDS-PAGE and refolded at a 1:1 ratio with *X. laevis* H2A. The H2A/H2B-Ub dimer was purified by size exclusion chromatography as described for unmodified H2A/H2B (Dyer et al., 2004). For fluorescently tagged histones, H2A(T120C) was labeled with maleimide derivatives of Cy3 or Cy3B prior to refolding as previously described (Shahian & Narlikar, 2012).

### Production of hexasomes and nucleosomes

Histone dimers (H2A/H2B), tetramers (H3/H4)_2_, and octamers (H3/H4/H2A/H2B) were refolded in equimolar ratios and purified by size exclusion chromatography as previously described (Dyer et al., 2004). Nucleosomes were generated by combining either the histone octamer or H2A/H2B dimer and H3/H4 tetramer (2:1 ratio) with DNA containing the Widom 601 sequence (Lowary & Widom, 1998). To favor hexasome formation, dimer and tetramer were combined in 1.2: 1 ratio. Reconstitution by salt dialysis was performed as described (Luger et al., 1999).

Addition of H2A/H2B dimer to hexasome was carried out within experimental reactions. H2A/H2B dimer (stored in 10 mM Tris (pH 7.8), 2 M NaCl, 1 mM EDTA, 5 mM 2-mercaptoethanol) was diluted roughly 10-30 fold to 6 µM in reaction buffer. Dimer dilutions were performed just prior to experiments. Hexasome was added to the reaction buffer first, followed by dimer in the indicated molar ratio. Dimer addition was allowed to proceed at room temperature for 2-3 minutes before additional reaction components were introduced. A similar pre-incubation step was carried out for nucleosome-containing reactions.

### Native gel sliding

Nucleosome sliding reactions were carried out as previously described with some minor adjustments (Eberharter et al., 2004; Patel et al., 2011). Briefly, 150 nM of fluorescently labeled nucleosome (or hexasome) and 50 nM Chd1 were diluted and combined in slide buffer (20 mM HEPES (pH 7.8), 100 mM KCl, 5 mM MgCl_2_, 0.1 mg・ml^−1^ BSA, 1 mM DTT, 5% sucrose (w/v)) at room temperature. Reactions were started with the addition of 2.5 mM ATP and at each timepoint, 1 µL of the reaction was added into into 24 µL of fresh quench buffer (20 mM HEPES (pH 7.8), 100 mM KCl, 0.1 mg・ml^−1^ BSA, 1 mM DTT, 5% sucrose (w/v), 5 mM EDTA, 125 ng/µL salmon sperm DNA (Invitrogen)) and placed on ice. To separate reaction products, 2.5 µL of the quenched timepoints were loaded onto 7% native polyacrylamide gels (60:1 acrylamide to bis-acrylamide) and electrophoresed (125 V) for 2 hours at 4˚C. Reaction products were observed by their fluorescent labels using a Typhoon 9410 variable mode imager (GE Healthcare).

### Histone mapping

Histone mapping experiments were conducted as previously described (Abbott et al., 2005). Briefly, nucleosomes made with fluorescently tagged DNA were labeled with ~220 uM p-azidophenacyl bromide (APB) on H2B S53C in the dark at room temperature for 3 hours before being quenched with DTT. Sliding reactions were assembled using 150 nM labeled nucleosome and 50 nM Chd1 in slide buffer (20 mM Tris Base (pH 7.8), 50 mM KCl, 5mM MgCl_2_, 5% sucrose (w/v), 0.1 mg・ml^−1^ BSA, 1 mM DTT). Before addition of ATP, a zero timepoint was collected. Sliding reactions were initiated with the addition of 2 mM ATP. At timepoints, 50 µL of the reaction was added to 100 µL of quench buffer (20 mM Tris Base (pH 7.8), 50 mM KCl, 5% sucrose (w/v), 0.1 mg・ml^−1^ BSA, 5 mM DTT, 5 mM EDTA, 150 ng/µL salmon sperm DNA) and placed on ice. The APB was crosslinked to the DNA by irradiating at 302 nm for 15 sec using a UV Transilluminator (VWR). Samples were denatured with 0.1% SDS and heat, and then subjected to phenol chloroform extraction and EtOH precipitation to remove uncrosslinked material. The crosslinked DNA was resuspended and cleaved with NaOH. The fragmented DNA was EtOH precipitated again before resuspending with formamide loading buffer and running it on an 8 M urea, 8% polyacrylamide (19:1 acrylamide:bis-acrylamide) sequencing gel. The samples were run for 1.25 hours at 65 W alongside a sequencing ladder of the nucleosomal DNA to allow precise identification of crosslink locations. Gels were imaged on a Typhoon 9410 variable mode imager (GE Healthcare) and analyzed using ImageJ (http://imagej.nih.gov/ij/).

### Exonuclease III digestion

Exonuclease III was carried out on nucleosomes and hexasomes with fluorescently labeled DNA. Samples containing nucleosome (100 nM) or hexasome (100 nM) preincubated for 2-3 minutes with 2-fold molar excess H2A/H2B dimer were incubated at room temperature for 10 minutes in reaction buffer consisting of 20 mM HEPES (pH 7.6), 50 mM KCl, 10 mM MgCl_2_, 5% sucrose (w/v), 0.1 mg・ml^−1^ BSA, and 1 mM DTT. For each sample condition, four 10 µL digestion reactions were made containing 0, 10, 40, and 160 units of Exonuclease III (New England Biolabs). After digesting for 5 minutes at room temperature, reactions were quenched by the addition of 40 µL of quench buffer (20 mM HEPES (pH 7.6), 50 mM KCl, 20 mM EDTA, 1.2% SDS) and placed on ice. DNA was isolated from the digestion reactions by adding an equal volume of phenol:chloroform:isoamyl alcohol (25:24:1), vortexing, centrifuging for 2 minutes, and removing the top (aqueous) layer to a new tube. To completely remove phenol, this step was repeated using chloroform:isoamyl alcohol (24:1). DNA was precipitated by adding 1.5 µL of 10 mg・ml^−1^ glycogen, 5 µL of 3 M sodium acetate and 250 µL 100% EtOH and then chilled at −80° C for > 20 minutes followed by centrifugation (21,130 rcf) for 30 minutes at 4° C. After a 70% EtOH wash and air drying of the pellet, samples were resuspended in 8 µL of formamide loading buffer and separated on urea sequencing gels as described for histone mapping.

### Single molecule FRET

Biotinylated and dye-labeled nucleosomes and hexasomes (alone or pre-incubated with an approximately twofold molar excess of unlabeled H2A/H2B dimer) were surface-immobilized on poly(ethylene glycol)-coated quartz microscope slides via a biotin-streptavidin linkage, as previously described (Blosser et al., 2009; Deindl et al., 2013). Immobilized samples were excited with a 532 nm Nd:YAG laser (CrystaLaser), and fluorescence emissions from Cy3 and Cy5 were detected using a prism-type TIRF microscope, filtered with a 550 nm long-pass filter (Chroma Technology), spectrally separated by a 635 nm dichroic mirror (Chroma Technology), and imaged onto the two halves of an Andor iXon Ultra 897 (512 × 512) CCD camera. The imaging buffer contained 12 mM HEPES, 40 mM Tris (pH 7.5), 60 mM KCl, 3 mM MgCl_2_, 0.32 mM EDTA, 10% glycerol, an oxygen scavenging system (800 µg ml^−1^ glucose oxidase, 40 µg ml^−1^ catalase, 10% glucose) to reduce photobleaching, 2 mM Trolox (Sigma) to suppress photoblinking of the dyes (Rasnik et al., 2006), and 0.1 mg・ml^−1^ BSA (Promega). Remodeling was induced by infusing the sample chamber with the imaging buffer containing 300 nM Chd1 remodeling enzyme and ATP using a syringe pump (Harvard Apparatus).

### Fluorescence Experiments

Static quenching of fluorescence (SQOF) experiments were carried out using Cy3-Cy3 or Cy3B-Dabcyl pairs. Reactions were monitored for 80N0 nucleosomes or hexasomes, with exit-side H2A C120 labeled with Cy3 or Cy3B and the zero-end of DNA labeled with Cy3 or Dabcyl (IDT). Sliding reactions were conducted with 10 nM nucleosome or 10 nM hexasome with 12 nM dimer (unless otherwise noted), 25 or 400 µM Chd1, and 25 µM ATP, 100 mM KCl, 20 mM HEPES (pH 7.5), 5 mM MgCl_2_, 100 µM EDTA, 5% sucrose w/v, 1 mM DTT, 0.2% Nonidet P-40, and 0.1 mg・ml^−1^ BSA at 25 °C.

Sliding reactions were monitored by either fluorometer or stopped-flow. Fluorometer experiments were conducted on a Fluorolog 3 fluorometer (Horiba) using a 2 mL reaction volume with a stir bar in the cuvette. First, 10 nM nucleosome or hexasome followed by H2A/H2B dimer was added to the cuvette and allowed to equilibrate for 2-3 minutes. Next, Chd1 was added, and after another brief equilibration, the sliding reaction was initiated with 25 µM ATP. Cy3 was excited at 510 nm and fluorescence was monitored at 565 nm using a 4 nm slit width and 1 second integration time. Stopped flow experiments were conducted on an SX20 stopped-flow (Applied Photophysics Limited) with nucleosome (or hexasome and dimer) and Chd1 in one syringe and ATP in the other. Cy3 was excited at 510 nm and emissions were monitored above 570 nm with a long-pass filter. Fluorescence signal was integrated over 0.01 seconds for the first ten seconds of the reaction and then 0.1 seconds for the remainder of the trace. Each progress curve is the average of 3-6 technical replicates. Progress curves were fit using the double exponential function, 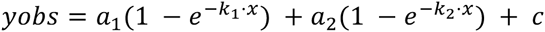, using Mathematica (Wolfram), where k_1_ and k_2_ are observed rates, a_1_ and a_2_ are corresponding amplitudes, and c is a constant.

## Acknowledgements

We thank Sarah Woodson for generously sharing equipment, Cynthia Wolberger and Mike Morgan for expression plasmids for Ubiquitin(G76C) and H2B(G120C), Sua Myong for suggesting use of Cy3B, and Ilana Nodelman for providing additional Chd1 protein and nucleosomes and hexasomes used for smFRET experiments. This work was funded by the National Institutes of Health (R01-GM084192 to G.D.B; T32-GM007231 to R.F.L.) and the KAW Foundation and Swedish Research Council (S.D.).

## Author contributions

R.F.L. and G.D.B. conceived of project, designed experiments, analyzed data, and wrote the paper; R.F.L. prepared reagents and performed experiments for hexasome and dimer addition characterization, Chd sliding, and Ubiquitin modification; A.S. and S.D. performed and analyzed single molecule experiments and edited the manuscript.

---

**Figure 1 - figure supplement 1. Hexasome orientation on the 601 is unaffected by the position of flanking DNA.**

**A.** Exonuclease III digestion performed as in Figure 1 except using 80-601-0 nucleosomes and hexasomes, where flanking DNA is on the opposite (left) side of the 601. More extensive digestion of hexasomes from the left of the 601 (lanes 5-8) indicates that hexasomes are missing H2A/H2B on the TA poor side of the 601 regardless of flanking DNA. Addition of two-fold molar excess H2A/H2B to hexasomes restored nucleosome digestion patterns (lanes 9-12). ExoIII digestions of 80-601-0 and 0-601-80 hexasomes and nucleosomes were performed in parallel, run on the same gel and are each representative of two independent experiments.

**B.** Histone mapping of 80-601-0 hexasomes and nucleosomes shows a loss of crosslinking from the TA poor side of the 601 (lane 26), consistent with 0-601-80 shown in Figure 1. Histone mapping of 80-601-0 constructs was performed alongside 0-601-80 constructs and run on the same gels. Results are representative of two or more independent experiments.

**Figure 5 - figure supplement 1. Remodeling saturates at 400 nM Chd1**

Nucleosomes formed by adding 12 nM dimer to 10 nM hexasomes were titrated with 10, 25, 50, 100, 200, 400, and 800 nM Chd1 and 25 µM ATP. Reactions were monitored by Cy3-Cy3 SQOF, and show that remodeling plateaued at 400 nM Chd1. The progress curves shown are representative of two independent Chd1 titrations using unmodified H2B (Wt-Wt). Similar results were observed for duplicate titrations using Ub-Wt, Wt-Ub, and Ub-Ub nucleosomes.

